# Lipid unsaturation properties govern the sensitivity of membranes to photo-induced oxidative stress

**DOI:** 10.1101/451591

**Authors:** A. Bour, S. G. Kruglik, M. Chabanon, P. Rangamani, N. Puff, S. Bonneau

## Abstract

Unsaturated lipid oxidation is a fundamental process involved in different aspects of cellular bioenergetics; dysregulation of lipid oxidation is often associated with cell aging and death. In order to study how lipid oxidation affects membrane biophysics, we used a chlorin photosensitizer to oxidize vesicles of various lipid compositions and degree of unsaturation in a controlled manner. We observed different shape transitions that can be interpreted as an increase in the area of the targeted membrane followed by a decrease. These area modifications induced by the chemical modification of the membrane upon oxidation, were followed in situ by Raman Tweezers Microspectroscopy (RTM). We found that the membrane area increase corresponds to the lipids peroxidation and is initiated by the delocalization of the targeted double bonds in the tails of the lipids. The subsequent decrease of membrane area can be explained by the formation of cleaved secondary products. As a result of these area changes, we observe vesicle permeabilization after a time lag that is characterized in relation with the level of unsaturation. The evolution of photosensitized vesicle radius was measured and yields an estimation of the mechanical changes of the membrane over oxidation time. The membrane is both weakened and permeabilized by the oxidation. Interestingly, the effect of unsaturation level on the dynamics of vesicles undergoing photooxidation is not trivial and thus carefully discussed. Our findings shed light on the fundamental dynamic mechanisms underlying the oxidation of lipid membranes, and highlight the role of unsaturations on their physical and chemical properties

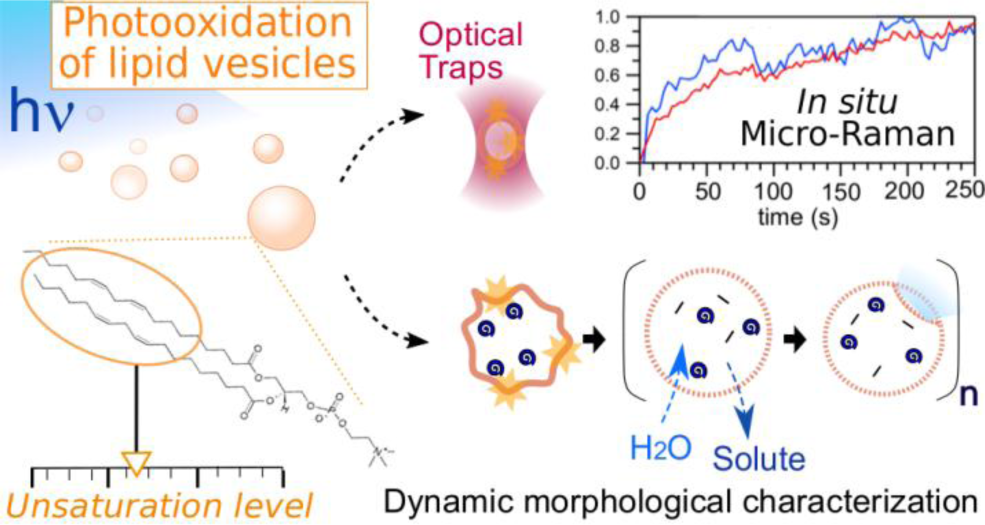

## INTRODUCTION

Aerobic metabolism in cells relies on the oxidation of organic compounds such as fatty acids, allowing cells to produce energy. However, the phospholipids that constitute cell membranes are also sensitive to oxidative stress, which can lead to membrane damage, aging and potentially cell death (1–3). The molecular processes responsible for the oxidation of unsaturated lipids involve the chemical modification of their double bounds and potential cleavage of the lipid chains (4, 5). Therefore, it is essential to consider the degree of unsaturation of fatty acids in order to understand the mechanisms allowing lipid membranes to withstand oxidative stress.

Biological membranes contain large amounts of mono- and polyunsaturated fatty acids, as well as traces of transition metals necessary for oxidative reactions. As they are constantly subjected to oxidation - in particular due to the respiratory chain - cellular membranes suffer from a variety of irreversible damages. Their oxidative state is defined as the balance between oxidant and antioxidant species, an imbalanced situation leading sequentially to stages of oxidative stress, cytotoxicity and pathological states (6, 7). Oxidative stress is known to be involved in cell aging and, if uncontrolled, in a variety of diseases including Parkinson’s and Alzheimer’s neurodegenerations and cancer (8–13). Oxidation-induced changes in membranes properties, such as fluidity and permeability (14–16), can result in abnormal and dysfunctional protein assemblies and in the leakage of vital molecules from organelles. Therefore, understanding membrane behavior with respect to their lipid unsaturation degree is an important biomedical challenge.

The first step of unsaturated lipid oxidation leads to the formation of hydroperoxides through a chain reaction ending when two lipid radicals react together (4, 5). Due to their polarity, the peroxidized hydrocarbon tails are thought to migrate to the lipid-water interface, which increases the molecule area and changes the packing parameter (17–19). The second step consists of further oxidation producing secondary products, including alcohols, ketones, alkanes, aldehydes and ethers (20, 21). These products are used as an index of lipid oxidation, mainly to estimate oxidation level of complex biological systems (22). The resulting cleaved lipids have a conical shape with a larger head to tail surface area ratio, and tend to increase the membrane permeability to solutes such as sucrose and glucose (23).

An effective and controlled way to induce oxidation processes in a system is to use photosensitizers to generate reactive oxygen species (ROS) upon light irradiation (24). Known as the photodynamic effect, the photochemical induction of oxidation is also involved in UV cytotoxicity and light-related ageing on tissues containing endogenous porphyrins (25). Moreover, together with the preferential retention of certain photosensitizers by tumors as compared to normal surrounding tissues (26), such photo-induced cytotoxicity has been applied as an anti-tumoral therapy known as the Photodynamic Therapy (PDT) (27).

Simplified models of biological membranes such as lipid vesicles are powerful tools for investigating how pho-to-induced oxidative species impact the phospholipid bilayer. Lipid vesicles exposed to photooxidation have been shown to exhibit a two-step behavior, corresponding to a sequential increase and decrease in membrane surface area. These modifications induce vesicle fluctuations and shape transitions (28–31), accompanied by modifications in the membrane viscosity (32), bending modulus (33), area expansion modulus (34), and permeability (28, 29, 35, 36). Similar effects have also been reported in polymersomes (37). Although correlations with chemical mechanisms have been proposed (30), the underlying dynamic chemical scenario remains largely unexplored due to a lack of *in situ* measurement during the oxidation process.

In this article, we investigate how lipid oxidation affects the dynamic behavior of artificial membrane models. We use Giant Unilamellar Vesicles (GUVs) made of controlled lipid composition with various degrees of unsaturation and expose them to photooxidation by light-activation of Chlorin e6 photosensitizer (Ce6, Fig. 1). Special attention is given to the number and position of lipid unsaturations. We first describe the time evolution of GUV morphology upon photooxidation and relate it to the membrane permeabilization. Then, we use Raman microspectroscopy combined with optical tweezers to follow *in situ* the membrane chemical modifications of vesicles in an aqueous environment. These results are discussed in relation to the observed mechanical and morphological changes.

**Figure 1.**
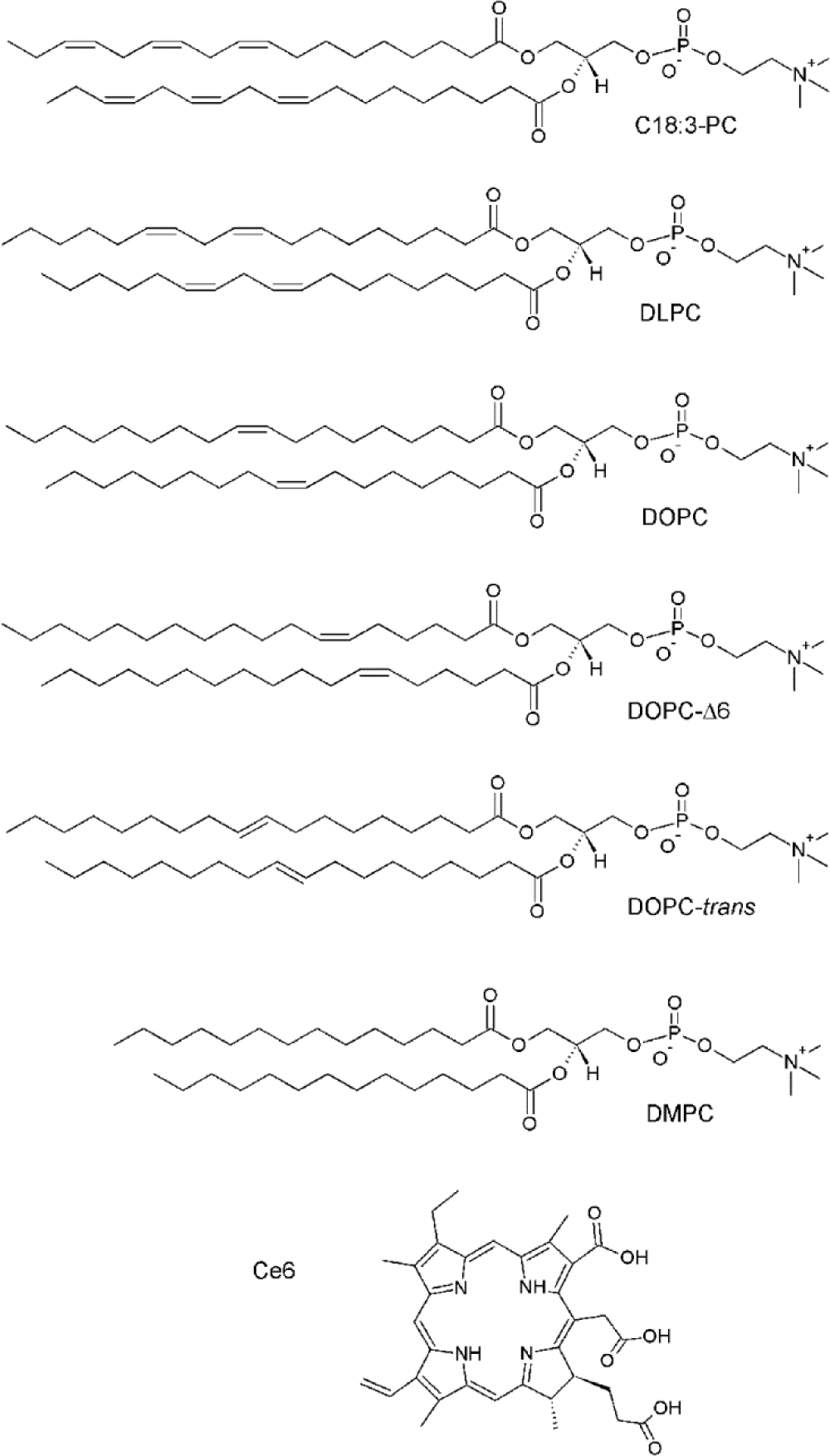
Formulae and abbreviations of the lipids composing the various GUVs, as well as of the photosensitizer Ce6.

## MATERIALS AND METHODS

### Chemicals

All chemicals were purchased from Sigma (USA), except the lipids (Fig.1) 1,2-dilinolenoyl-sn-glycero-3-phosphatidylcholine (C18:3-PC), 1,2-dilinoleoyl-sn-glycero-3-phosphatidylcholine (C18:2, DLPC), 1,2-dioleoyl-sn-glycero-3-phosphatidylcholine (C18:1, DOPC), 1,2-dipetroselenoyl-sn-glycero-3-phosphatidylcholine (DOPC-∆6), 1,2-dielaidoyl-sn-glycero-3-phosphatidylcholine (DOPC-trans), 1,2-dimyristoyl-sn-glycero-3-phosphatidylcholine (C14:0, DMPC) from Avanti Polar Lipids (USA), and chlorin e6 from Frontier Scientific (USA). The lipids were handled very carefully, in vaccuum or under nitrogen atmosphere, when possible. For all lipids, the absence of oxidation is measured by Raman (spectra at t=0). Chlorin stock solution (5 mM) was prepared in ethanol and kept at –18 °C. The experimental Ce6 aqueous solutions were prepared and used without delay and handled in the dark. The osmolarity of the solutions was checked with an osmometer (Löser Messtechnik, Germany).

### Vesicle formation

The lipids (C18:3-PC, DLPC, DOPC, DOPC-trans or DOPC-∆6) were solubilized in chloroform. For the determination of the binding constant between lipid membranes and Ce6, Large Unilamellar Vesicles (LUVs) were prepared by extrusion method (38). After evaporation of chloroform, lipids were dispersed in PBS buffer by vortexing. Using an extruder device (Avanti Polar Lipids, USA), the liposome suspension was extruded 8–10 times through a polycarbonate membrane filter (Poretics, Livermore, CA) with pores of 200 nm.

The electroplating method (39) was used to form giant unilamellar vesicles (GUVs) of a diameters of 20 to 30 μm. Lipid mixtures in chloroform were deposited on ITO-covered glass plates, and chloroform was evaporated in vacuum. A formation chamber was then made from two such glass plates and a Teflon spacer of 4 mm. The chamber was then filled with a solution of 300 mM sucrose and HEPES buffer solution (2 ml, pH 7.8, HEPES 0.5 mM). An AC field of 1 Volt and 8 Hz was applied between the plates for 4 hours. For experiments, the GUVs were mixed with a ~300 mM glucose solution made in the same buffer. The solution carefully adjusted ~5 mOsm/L higher than the external sucrose solution to limit the strain of the membrane. The density difference between sucrose and glucose caused the GUVs to sediment to the bottom of the chamber. The difference in optical index between sucrose inside and glucose outside the vesicle allowed phase contrast microscopy observations.

### Measurements of the Ce6 affinity to membranes

For the steady-state study of the interaction between the Ce6 lipid membranes of various compositions fluorescence spectra were measured with an Aminco Bowman Series 2 spectrofluorimeter (Edison, NJ, USA). All experiments were done in triplicates. LUVs solutions were prepared at different concentrations of lipids and 10 μL of 10 μM Ce6 solution were added to 2 mL of each vesicles preparation. The fluorescence spectra were recorded 2 minutes after the sample preparation. Data thus obtained were analyzed as described elsewhere (28, 28, 40, 41). The fluorescence intensity at a wavelength corresponding to the maximum of fluorescence emission of Ce6 incorporated into the membrane was plotted as a function of the lipids concentration. The binding constant, K_B_, was derived according to the relationship :

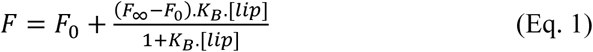

where F_0_, F_∞_ and F are the fluorescence intensities corresponding to zero, total and intermediate incorporation of Ce6 into vesicles respectively. Lipids molecules are in large excess and the bilayer is far from being saturated with Ce6. Therefore, it was assumed that the membrane properties were independent of the number of bound Ce6 molecules and [lip] was equal to the total lipid concentration.

Partition experiments were carried with an excitation wavelength at 410 nm. Upon the addition of vesicles, the intensity of the fluorescence emission increases significantly and shifts from 660 to 668 nm. Typical fluorescence emission spectra of the photosensitizer in solution and in the presence of various DLPC lipid concentrations are shown in Fig. 2 (inset). These spectral changes are characteristic of the transfer of the chlorin from an aqueous to a hydrophobic environment (28, 36, 40, 42). The plot of the fluorescence intensity at 668 nm versus the lipid concentration shows a saturation profile (see Fig. 2). The binding constant value, K_B_, was determined by fitting the experimental data with Eq. 1 and is reported in Table 1. The obtained values are in good agreement with previous data obtained (28, 36) and depend of the membrane composition. Such an effect has been previously observed for the interactions of photosensitizers with various membrane (26, 28, 37, 40, 42).

**Table 1.**
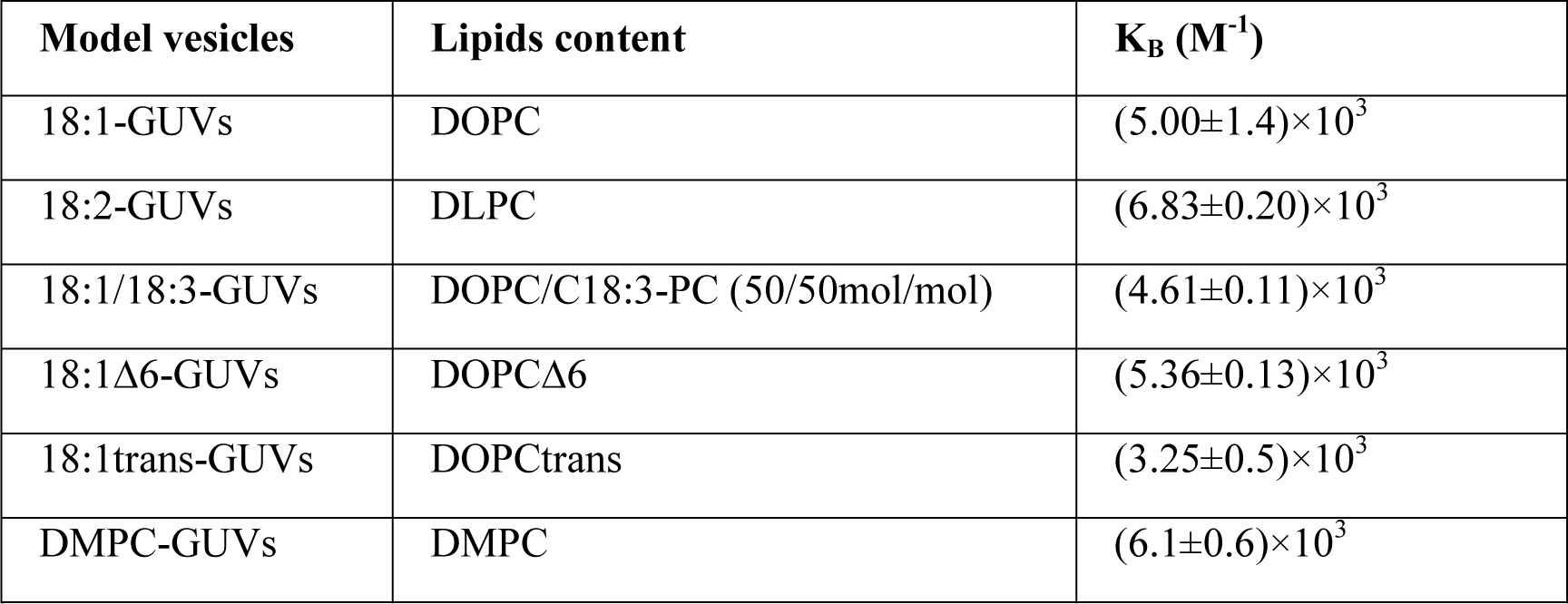
Lipid content and Ce6 binding constants for each type of model vesicle

| Model vesicles | Lipids content | $K_B$ ($M^{-1}$) |
| --- | --- | --- |
| 18:1-GUVs | DOPC | $(5.00 \pm 1.4) \times 10^3$ |
| 18:2-GUVs | DLPC | $(6.83 \pm 0.20) \times 10^3$ |
| 18:1/18:3-GUVs | DOPC/C18:3-PC (50/50mol/mol) | $(4.61 \pm 0.11) \times 10^3$ |
| 18:1Δ6-GUVs | DOPCΔ6 | $(5.36 \pm 0.13) \times 10^3$ |
| 18:1trans-GUVs | DOPCtrans | $(3.25 \pm 0.5) \times 10^3$ |
| DMPC-GUVs | DMPC | $(6.1 \pm 0.6) \times 10^3$ |

**Figure 2.**
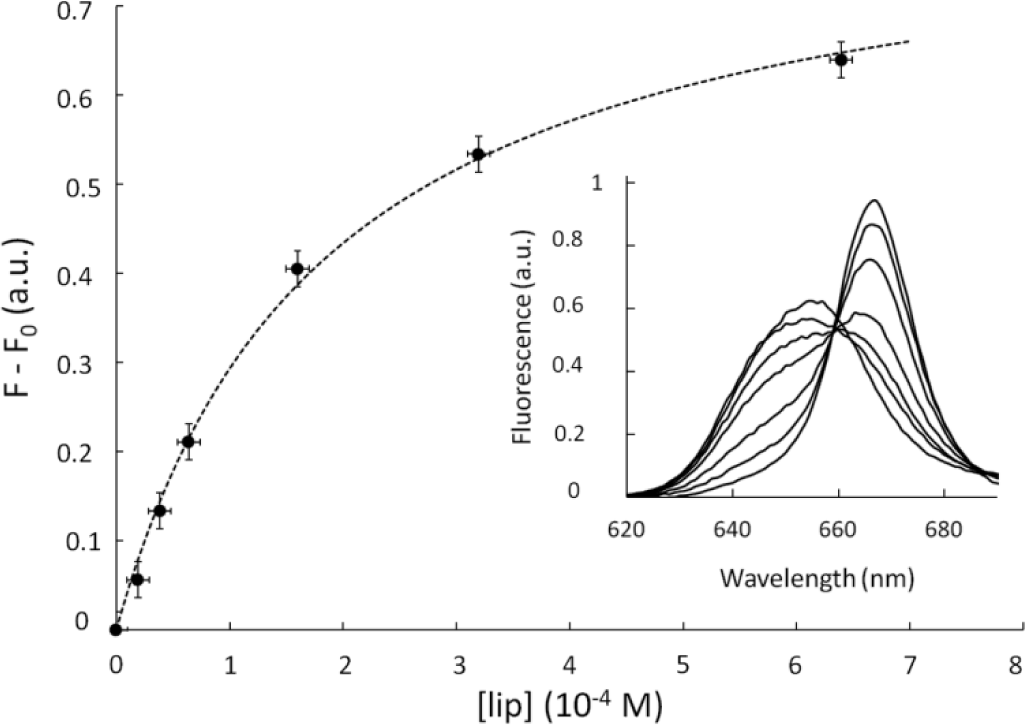
Incorporation of Ce6 within lipid membranes. Evolution of the Ce6 fluorescence emission upon incorporation into vesicles (here, made with pure DLPC; Ce6 concentration is 5×10^-8^ M). The excitation wavelength is 410 nm and the fluorescence intensity is recorded at 668 nm. Eq. 1 is used to fit experimental data and yields the binding constant K_B_. The inset shows the red shift of Ce6 fluorescence peak upon incorporation within the membrane.

### Labeling of GUVs

Giant Vesicles were labeled with Ce6 as previously reported (28, 29). The Ce6 concentration used was calculated according to the K_B_ value determined beforehand for each lipid mixture to ensure an identical membrane labeling. The homogeneous distribution of the fluorescent Ce6 molecules within the vesicle membranes was assessed by fluorescence microscopy.

### Observation and illumination

GUVs were observed under an inverted microscope (Nikon Eclipse TE 300 DV) equipped with a high numerical aperture phase oil objective (CFI Plan apochromat DM 60 n.a: 1.4, Nikon, France). Illumination was provided by a 120 W Metal halide lamps with a 405±10 nm bandpass filter. Neutral density filters (NDx8) were used to reduce illumination level. GUVs were simultaneously imaged in bright-field using the halogen lamp of the microscope and the fluorescence of Ce6 was cut off by using an appropriate filter. Image acquisition (100 ms integration time) was performed with a CCD camera (Neo sCMOS, Andor Technology). The acquisition, processing and image analysis were performed with NIS-Element provided by Nikon and Image J (NIH, USA). For each membrane composition, the images of at least twenty vesicles from three sample preparations were analyzed separately. The intra-trial variability is reported in the supporting information. When photo-oxidizing the membrane, illumination was kept constant from the initial time.

### Chemical scenario

To monitor the chemistry of lipid oxidation, Raman spectra of corresponding lipid vesicles in the presence of photosensitizer Ce6 were recorded using home-built Raman Tweezers Microspectroscopy (RTM) setup described elsewhere (43). Briefly, RTM technique combines optical trapping (44) with Raman probing: the same laser radiation is used for both optical trapping of the vesicles and excitation of Raman scattering from the vesicles’ constituent biomolecules. The light at 780 nm is provided by a continuous-wave Ti:Sapphire laser (Spectra Physics, model 3900S) pumped by argon-ion laser (Spectra Physics Stabilite 2017). The laser beam is focused by a water-immersion infinity-corrected objective (Olympus LUMFL 60X, NA 1.1) brought into contact with a droplet (~100 µL) of water buffer containing the vesicles of interest. The laser power was adjusted to about 70 mW inside the sample suspension. LUVs concentration was adjusted to about 100 µg/mL in lipids and to about 0.1% in Ce6. Our RTM setup employs upright microscope configuration and the same objective is used to collect Raman signal in a back-scattering geometry and to deliver it into a spectrograph (Acton SpectraPro 2550i) coupled with a deep-depleted back-illuminated NIR CCD (Princeton Instruments SPEC-10 400BR/LN). Raman light is focused onto the spectrograph’s entrance slit (semi-confocal configuration, slit width 50 µm) by an achromatic lens with f=75 mm. The Raman signal is spectrally separated by 2 Semrock RazorEdge long-pass filters (grade ‘‘U’’): one (blocking filter) with the edge wavelength of 780 nm is placed normally to the optical beam just before the focusing lens; another one with the edge wavelength of 830 nm is used as a dichroic beam splitter at an angle of incidence 45°: it reflects laser light at 780 nm and transmits all the wavelengths longer 785 nm. Spectral resolution in all Raman experiments was about 5 cm^-1^. Frequency calibration was performed using Raman lines of toluene with ±2 cm^-1^ absolute accuracy and relative frequency position accuracy better than ±1 cm^-1^.

All experiments were done in triplicates. Raman spectra were continuously acquired every 3 seconds within 10-minutes intervals using WinSpec software; further data treatment was performed using IgorPro for Windows software. The informative signal from the vesicles in the sample volume arises as additional spectral features appearing on top of water buffer Raman spectrum, as soon as the vesicles are trapped at the focus of the laser beam. Simultaneously with the event of vesicles trapping, the sample droplet was illuminated by weak light at 405 nm from the microscope white-light illumination system passing through the band-pass interference filter (Andover Corporation 405±5 nm), in order to initiate the photosensitizing effect. The details of raw Raman spectra treatment were described in detail elsewhere (43). In this study, each spectra is the mean of three acquisitions. For Raman kinetics evaluation based on spectral band areas, an additional step of spectra treatment was introduced, consisting in automatic background correction using linear functions and spectra normalization on ν(C-N+) stretch of polar heads.

## RESULTS AND DISCUSSION

### Preparation of the photooxidable vesicles

In order to investigate the effect of lipid unsaturation on membranes undergoing photooxidation, we use electroformed GUVs made of five distinct compositions. To vary the level of membrane unsaturation we considered (i) 18:1-GUVs, made of pure mono-unsaturated lipids DOPC, (ii) 18:2-GUVs, composed of pure polyunsaturated lipids DLPC and (iii) 18:1/18:3-GUVs, a mixture of C18:3-PC and DOPC at 50/50 molar ratio that have a mean level of unsaturation similar to DLPC. Then, to address the role of the geometry of the unsaturation, we used (iv) 18:1trans-GUVs, composed of DOPC-trans lipids and (v) 18:1∆6-GUVs, made of DOPC-∆6 lipids. All unsaturated lipids used here were C18 phosphatidylcholine, in order to address the effects of the unsaturation of the lipids tails and not that of their head group, nature or length. Control experiments were carried using GUVs made of the saturated lipid DMPC. In all cases, the photosensitizer Ce6 was added to the GUVs membranes to enable the photo-control of the membrane oxidation. Beforehand, partition experiments were carried out to quantify the propensity of the Ce6 to insert within the various membranes and determine the suitable Ce6 concentration for each membrane composition (see binding constants in Table 1).

The GUVs were prepared in a solution of sucrose and diluted in a glucose solution to produce a difference in optical index between the inner sucrose and outer glucose solutions, facilitating observations and measurements by phase contrast microscopy. GUVs were observed by bright field optical microscopy and photooxidation was induced by illumination with a 120 W Metal halide lamp with a 405±10 nm bandpass filter.

### Morphological evolution of GUVs under photooxidation

In the absence of irradiation, all GUVs appeared spherical, although slightly floppy due to the small osmolarity mismatch between the inner and outer solutions (5 mOsm/L, see Materials and Methods for details). As seen from Fig. 3, upon illumination, the Ce6-labeled GUVs exhibited a dynamic morphology characterized by two consecutive stages: first a transient shape destabilization with the appearance of outwards buds or membrane invaginations (Stage 1), followed by a recovery of a spherical shaped accompanied by a progressive decrease in optical contrast and in certain cases a reduction in vesicle size (Stage 2). In comparison, control vesicles either containing no Ce6 or lacking unsaturations (DMPC-GUVs labeled with Ce6), showed no shape changes upon illumination, confirming that the observed dynamics is solely induced by the oxidation of unsaturated lipids through the photosensitizer Ce6.

**Figure 3.**
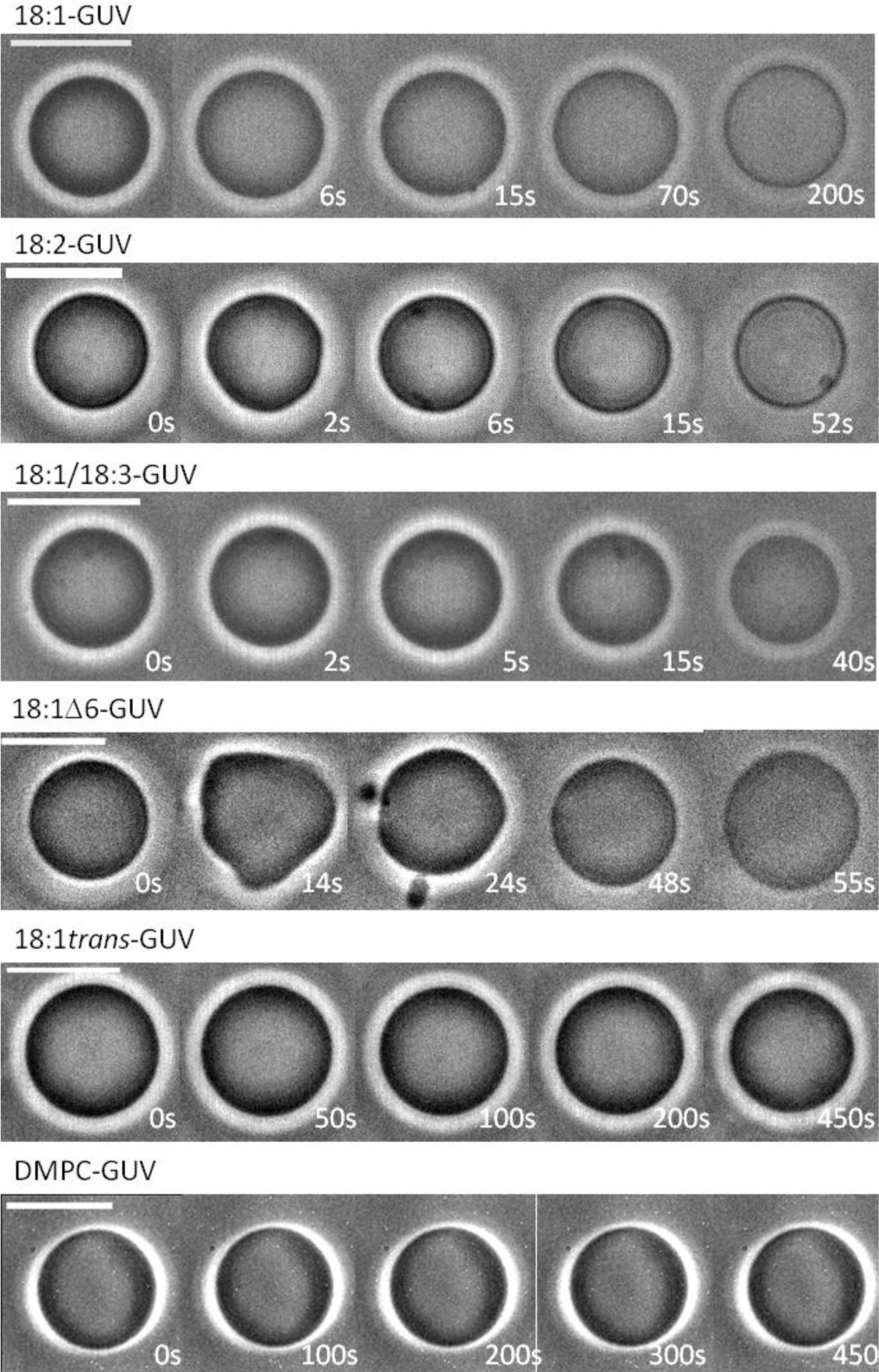
Effect of lipid unsaturation on the morphological evolution of GUVs under photooxidation: Morphological changes of Ce6-labeled GUVs of various compositions exposed to photooxidation. Shortly after irradiation, GUVs enter a first stage characterized by an increase of membrane surface area leading to shape destabilization. Then, during a second stage, a decrease of membrane area allows the GUVs to recover a spherical shape, concomitant with a loss of contrast. Irradiation starts at time t=0s and time scale is adjusted according to the duration of the processes i.e. the composition of the vesicles. The figure presents images of vesicles of comparable radii (~5 µm); the intra-trial variability, related to the system size, is presented in the supporting information. Scale bar = 10 µm.

The shape transitions observed in Stage 1 are consistent with reported dynamics of lipid vesicles of various composition containing different photosensitizers such as chlorins, fluorescent probes or methylene blue (28–31, 34). The shape destabilization is attributed to a transient increase in membrane area, possibly coupled to a softening of the lipid bilayer mechanical properties. Most importantly, we find that the kinetics of shape transition is strongly dependent on the unsaturation properties of the membrane lipids. Although the duration of Stage 1 is of the order of 10s for both 18:1-GUVs and 18:2-GUVs, it is decreased to about 2 seconds for 18:1/18:3-GUVs. For 18:1∆6-GUVs whose double bond is nearest the lipid head, Stage 1 lasts for up to 30s and presents shape fluctuations of higher amplitudes. In contrast, 18:1trans-GUVs seem unaffected and retain their spherical shape, initial radius and optical contrast for all the duration of the experiment (450s).

### Pulsatile dynamics of photooxidized vesicles

In order to quantitatively compare the effects of lipid unsaturation on the dynamics of GUVs undergoing photooxidation, we recorded for each vesicle composition the time evolution of (i) their optical contrast and (ii) their radius. As seen Fig. 4, all photosensitive GUVs showed a dramatic decrease in optical contrast during Stage 2. Given that the optical contrast is the result of the difference in sugar composition between the interior (sucrose) and exterior (glucose) of the GUVs, the decrease in contrasts is interpreted as a combined effect of the dilution and leak-out of encapsulated sucrose due to a progressive permeabilization of the lipid membrane.

**Figure 4.**
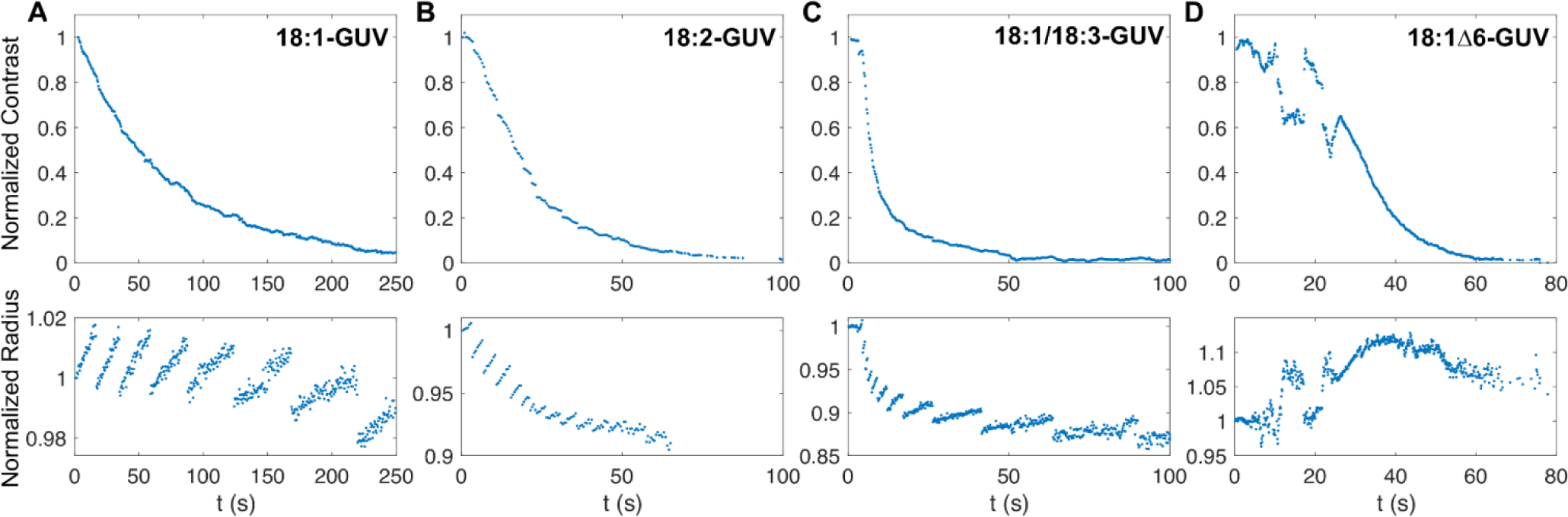
Time evolution of optical contrast and radius of GUVs undergoing photooxidation. (Top) Evolution the contrast between the inside and the outside of the oxidized vesicles. The decay is the result of a decrease in concentration difference between the GUVs inner sucrose solution and the surrounding glucose solution. (Bottom) Vesicle radii exhibit cycles of progressive increases followed by abrupt drops, characteristic of GUVs swell-burst cycles. The results presented are those of the same vesicles as those in Figure 3.

Strikingly, the vesicle radii exhibited an overall decrease, marked by pulsatile behavior of progressive swelling and abrupt drop for 18:1-GUVs, 18:2-GUVs and 18:1/18:3-GUVs. This dynamics is similar to those of lipid GUVs undergoing swell-burst cycles induced by osmotic stress (45–47) or subject to surfactants (48, 49). Based on these observations, we propose the following mechanism for the GUVs dynamics (Fig. 5). Upon illumination, the Ce6 labeled GUVs generate reactive oxygen species (ROS), that lead to oxidative reactions of the unsaturated lipid tails. At short time scales (~10s), the oxidation produces an increase in membrane area resulting in shape destabilization of the initially spherical vesicles (Stage 1) (30). At longer time scales, the surface area of photooxidized membranes progressively reduces, resulting in a recovery of the spherical shape followed by a decrease in the GUV volume (Stage 2). Concomitantly, oxidative products are released from the membrane both in the intra-vesicular space and in the surrounding bath. However, because the volume of the bath is much larger than the vesicle volume, a positive concentration difference builds up between the inside and the outside of the GUV. Since the permeability of the membrane to oxidative product is negligible when compared to its permeability to water, the concentration difference induces an hypotonic stress driving an inflow of water through the membrane and the progressive swelling of the GUV. Lipid membranes are elastic materials able to withstand about 4% surface strain before rupturing (50). Once this critical strained is reached, the GUV opens a micrometer sized pore through which the inner solution leaks out, relaxing the internal pressure and decreasing in turn the vesicle volume and membrane tension. This in turn allows for the pore line tension – induced by the hydrophobic lipid mismatch at the pore edge – to drive the resealing of the pore (47). Such swelling and bursting sequence is repeated in successive cycles as long as oxidative species are released from the membrane. The 18:1∆6-GUVs showed a slightly different behavior, with no clear swell-burst cycles, although these may happen with amplitudes smaller than our setup can detect.

**Figure 5.**
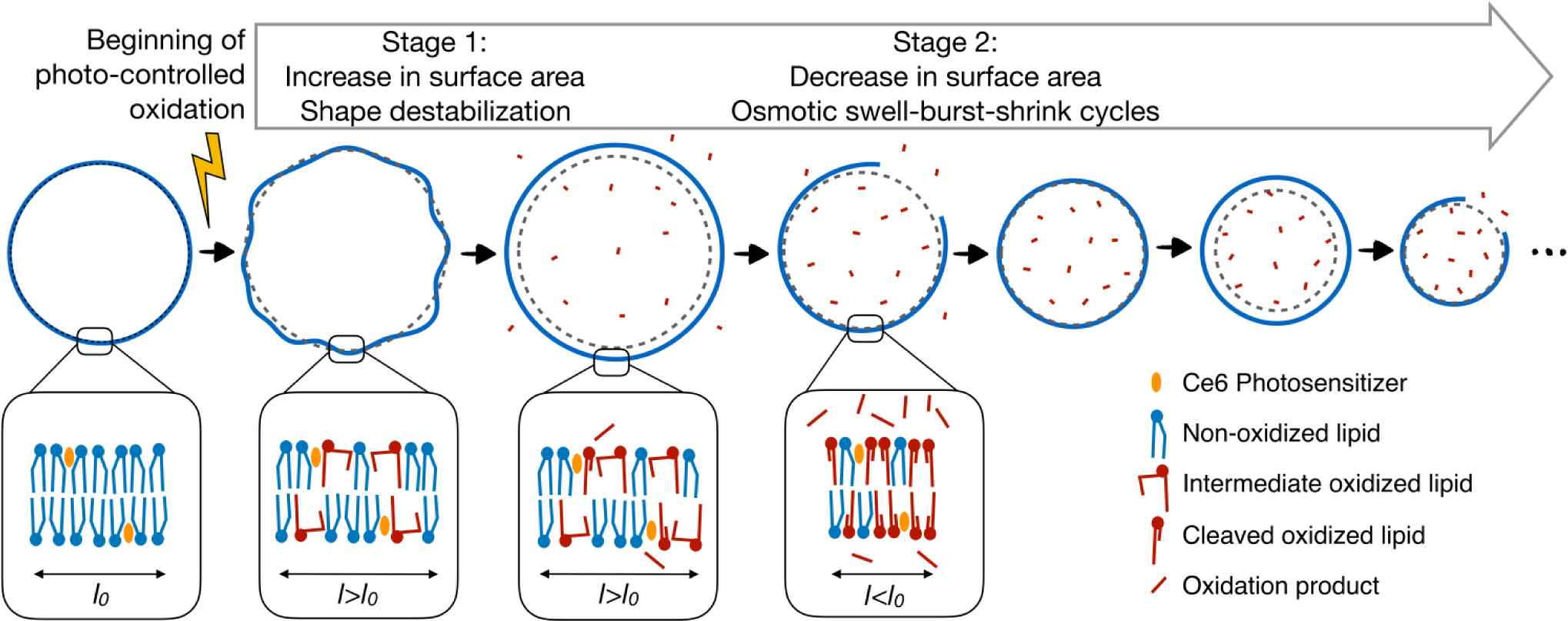
Sketch of the proposed mechanism involved in the dynamics of the vesicles over oxidation.

To confirm that the swell-burst cycles occurred in 18:1-GUVs, 18:2-GUVs and 18:1/18:3-GUVs, we tracked the vesicles with rapid video-microscopy (100 Hz) in search of a transient pore opening. As shown in Fig. 6, we observed short lived pore opening up to few microns in radius for about 50 ms, thus confirming the proposed swell-burst mechanism. We then calculated for each cycle the maximum area strain (Fig. 7, A-D). For the GUVs for which pore openings have been demonstrated, this corresponds to the critical strain ε_c_ at which membrane bursts. For 18:1-GUVs and 18:2-GUVs, the maximum strain decreases with each cycle, reaching a final value about four times lower than at the beginning of Stage 2 : ε_c_ is decreased respectively for 18:1-GUVs and 18:2-GUVs from (6.61±2.07) and (3.5±1.00)% down to (1.48±0.47) and (1.3±0.5)%. This decrease suggests a weakening of the membranes undergoing oxidation. In contrast, for 18:1/18:3-GUVs, the maximum strain stays around 3%, with a slight increase within the range of the statistical errors.

**Figure 6.**
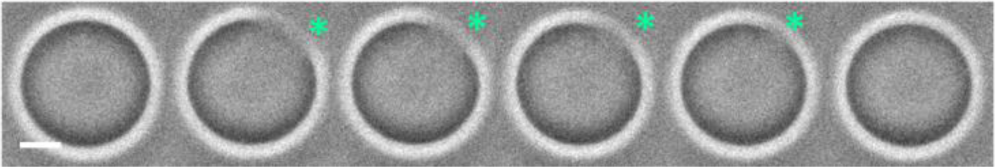
Time evolution of a transient pore opening in a 18:2-GUV under photooxidation. The pore is marked with an asterisk. Frequency of image acquisition: 100 Hz. Scale bar: 10 µm.

**Figure 7.**
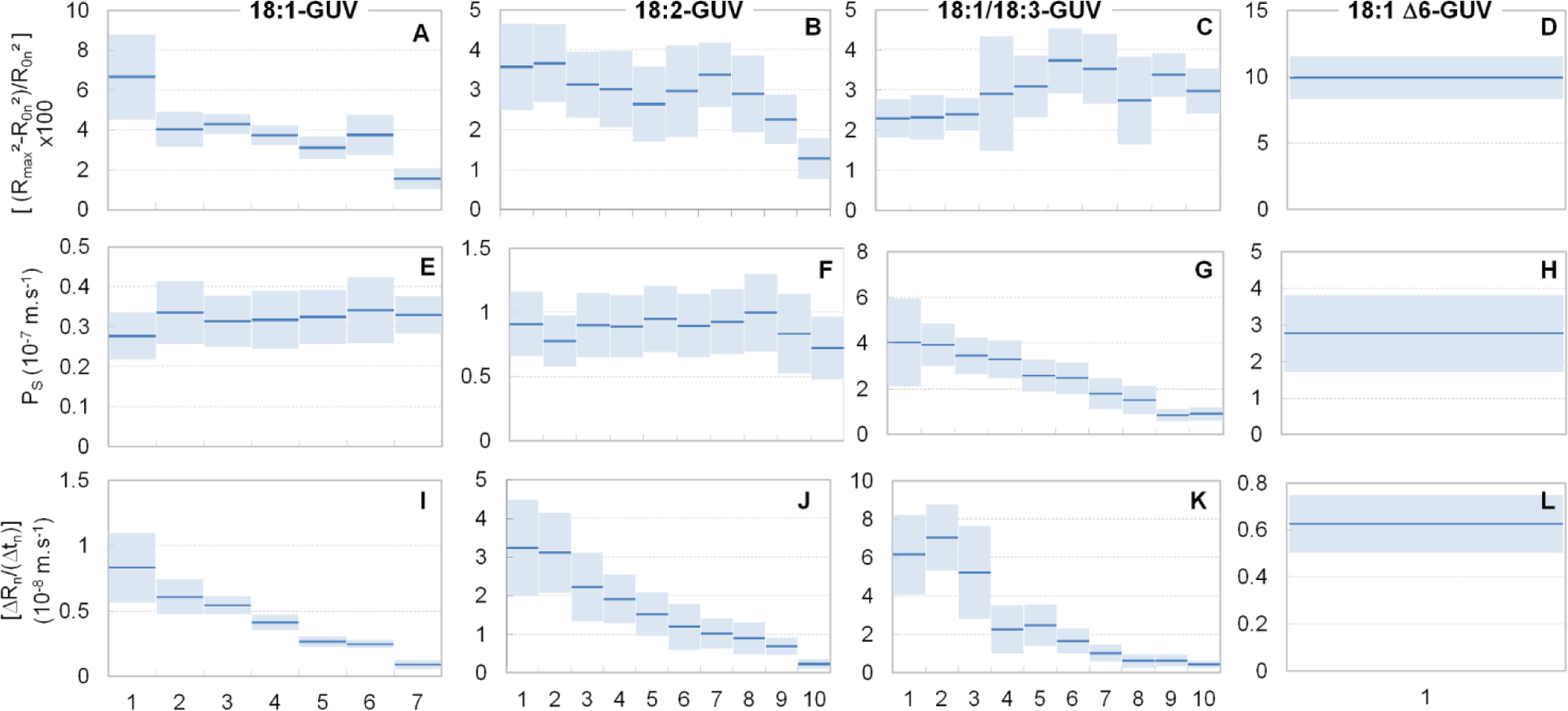
Evolution of physical parameters of GUVs undergoing photooxidation as a function of the swell-burst cycle number. (A-D) Evolution of the membrane critical strain, 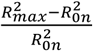. (E-H) Permeability to sucrose, *Ps*. (I-L) Progressive decrease of the water influx related term of Eq. 4, 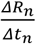. The 18:1Δ6-GUVs are analyzed as if they were undergoing a unique cycle. Dark lines are the mean of 20 vesicles per compositions, error bars are ± standard deviations.

### Contribution of membrane permeabilization to GUV contrast decay

From our observations of transient pores (Fig. 6), we did not find any significant drop in vesicle contrast with pore opening, suggesting that the short times of the pore opening events do not allow diffusion through the pore to affect the concentration imbalance in sucrose and glucose. Indeed, convection driven leak-out alone does not change the concentration of sucrose inside the vesicle (47). Therefore the main mechanisms leading to the progressive loss in contrast observed in Fig. 3 are the dilution of the inner content by the osmotic influx of water, as well as the permeabilization of the membrane to sucrose upon oxidation, in accordance with the packing defect and/or pre-pore opening scenarios, where the evolution of hydrophobic defects of the membrane involve local increase in spacing of lipid molecules (28, 29, 35, 51, 52).

In order to evaluate the permeabilization of the lipid membrane by photooxidation, we propose a mathematical model of the swell-burst process (see Supporting Information for full derivation). Based on this framework, we estimate the time evolution of the intra-vesicular sucrose concentration (*c_s_*) during a given cycle (*n*):

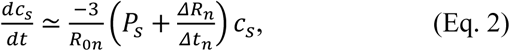

where *R_0n_* is the initial vesicle radius of the *n*^th^ cycle, *P_s_* is the membrane permeability to sucrose, *∆R_n_* is the total amplitude of the vesicle radius during that particular cycle and *∆t_n_* is the cycle period. This equation states that the sucrose concentration in the vesicle can decrease either by permeating through the membrane (first term in parenthesis) or by dilution when the influx of water increases the vesicle volume (second term in the parenthesis). Assuming that the membrane permeability to sucrose is constant within the period of a cycle, the solution of Eq. 2 is an exponential decay of the form

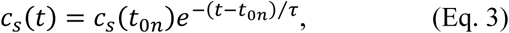

where *t_0n_* is the time at which the cycle starts and *τ* is the characteristic decay time of the sucrose concentration defined as

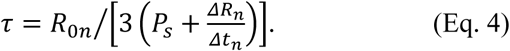

The two terms in the denominator of Eq. 4 reflect the two processes involved in the progressive loss of contrast of the vesicles, i.e. respectively the permeabilization of the membrane to the sucrose and the osmotic influx of water. For each GUVs cycle, the measured quantities are *∆R_n_*, *∆t_n_*, *R_0n_* and *t_0n_*. Therefore, assuming that the optical contrast is proportional to the sucrose concentration in the GUVs, one can determine *Ps*, the membrane permeability to sucrose, by obtaining τ from the fit of Eq. 3 to the contrast decay during a given cycle. *ΔR_n_/Δt_n_* is directly derived from the radii measurements (Fig. 3). We observed that the magnitude of this term represents less than 20% of *Ps* for 85% of the cycles and that its contribution decreases along the permeabilization (see Fig. 7, E-L). Therefore, the loss of contrast observed in Fig. 4 is essentially governed by the permeabilization of the membrane to the sucrose.

Moreover, except for 18:1/18:3-GUVs, we did not evidence any important variation in membrane permeability over the oxidation time (Fig. 7), suggesting that the oxidation of the membrane induces an increase of its permeability to sucrose in an abrupt manner, by three order of magnitude, a native phospholipid membrane being almost impermeable to sucrose (Ps* = ~10^-12^-10^-10^ m/s). Interestingly, such an observation suggests a threshold effect between two distinct permeability regimes for native and oxidized membranes, regardless of their oxidation degree and the level of oxidized products (16). So, the GUVs contrast decay is characterized by two parameters: *t_0_*, the time of onset of contrast decay and *P_s_*, the permeability to sucrose of oxidized membranes, on which the characteristic decay time essentially depends. The measured values of *t_0_* for 18:1-GUVs and 18:2-GUVs (9.4±4.5 and 10.1±5.1 seconds, respectively) do not show significant differences. This indicates that the onset of the membrane permeabilization is independent of the number of double-bond in the lipid tail. However, the *t_0_* obtained for 18:1∆6-GUVs is 28.4±5.5 seconds, three fold higher than 18:1-GUVs and 18:2-GUVs, suggesting that the position of the unsaturation determines the initial permeability resistance to photooxidation. For 18:1/18:3-GUVs, *t_0_* value is 2.6±0.8 seconds, four fold smaller than for 18:1-GUVs, revealing the weak stability of this membrane upon oxidation.

In contrast, the value of *P_s_* (Fig. 7, E-H) of oxidized membranes increases with the number of unsaturation from (0.32±0.02)x10^-7^ m.s^-1^ for 18:1-GUVs to (0.87±0.08) 10^-7^ m.s^-1^ for 18:2-GUVs. This behavior indicates a direct correlation between the number of double bounds and the oxidized membrane permeability. For 18:1/18:3-GUV, after the abrupt increase in the value of *Ps* compared to that of non-oxidized membranes, we observe throughout Stage 2 a decrease of permeability to sucrose from (3.87±0.89)x10^-7^ m.s^-1^ to (0.88±0.24)x10^-7^ m.s^-1^, a value of same order of magnitude than for 18:2-GUVs but that remains larger than for unoxidized membrane. This suggests a reorganization in the lipid membrane, in parallel with its oxidation, possibly involving changes of the structural or dynamics properties such as lipid packing and order (53, 54). The fastest loss of contrast occurs for the 18:1Δ6-GUV, with a *P_s_* value of (1.79±0.81)x10^-7^ m.s^-1^, showing an oxidation dependence at the position of the unsaturation.

### Chemical dynamics of membrane photooxidation

The above analysis gives valuable insight into the mechanical and transport properties of the various GUVs undergoing photooxidation. Yet, the underlying chemical scenario is largely unexplored. Indeed, because reaction intermediates are very short-lived, following the photochemistry involved in membrane alteration *in situ* is challenging. We used Raman Tweezers Microspectroscopy (RTM) to monitor simultaneously the kinetics of photochemistry involved in Large Unilamellar Vesicles (LUV) oxidation and the morphological response of the vesicles. The main advantage of this technique is that it allows us to follow the dynamics of oxidative cascade *in situ*. Vesicles labeled with Ce6 were trapped by a strong NIR light at 780 nm and simultaneously irradiated by a weak near-UV light at 405 nm, in order to initiate the photoinduced oxidation. The RTM method has been described elsewhere (43, 44) and is detailed in the Supporting Information. The changes in the Raman spectra of the vesicles lipids were recorded over 10 minutes, with an accumulation time of 3 seconds per spectrum effectively determining the time resolution of our RTM kinetic measurements. Raman spectra of Ce6 labeled LUVs composed of unsaturated lipids exhibited bands corresponding to the vibrations of the hydrocarbon chains with some contributions of the polar head groups (55, 56). The assignment of the characteristic Raman bands in spectra of lipid vesicles under study are presented in Table S1. The characteristic photo-induced changes vary depending on the degree of unsaturation. In general, the following processes have been detected and quantified: peroxide formation, through the appearance of transient Raman band at 840-880 cm^-1^; C=C bond isomerization, via the appearance of characteristic derivative-like feature at 1650-1675 cm^-1^ in difference Raman spectra; conjugation of C=C bonds, through the appearance of extra intensity around 1654 cm^-1^; and a cyclization reaction, via the appearance of strong 6-membered ring bands at 1000, 1031 and 1596 cm^-1^. Note that no photoinduced changes were observed for 18:1*trans*-LUV.

As shown in Fig. 8, all of the examined vesicles exhibit changes of the bands which were involved in C=C loss or isomerization. All photooxidized lipids also displayed an apparition of a band in the 820-860 cm^-1^ region corresponding to the peroxide apparition signed by an increase of the O-OH stretching mode. The oxidation process initiates through the delocalization of the double bonds (C=C), which move along the fatty chain. This results in the formation of *trans* and conjugated double bonds, for the mono- and poly-unsaturated lipids, respectively. From the kinetics point of view, the rate of C=C attack is directly correlated with the number of unsaturation. Moreover, the isomerization of 18:1Δ6-LUV is three fold faster than for 18:1-LUV (see supporting information: the equalization of the *cis* and *trans* C=C bands intensities takes 20 seconds for DOPCΔ6, versus 54 seconds for DOPC). This suggests an important role of the C=C bond position. In parallel to this initial step of double bonds delocalization, the peroxidation reaction with oxygen develops, leading to the primary products, the hydroperoxides. The formation of these transient compounds is visible in all spectra – except for 18:1trans-GUVs, where no changes where evidenced. This suggests that the initial delocalization step is necessary for the reaction with oxygen, as attested by the absence of peroxidation of the 18:1*trans*-vesicles where the peak at 1670 cm^-1^ remains stable over irradiation time.

**Figure 8.**
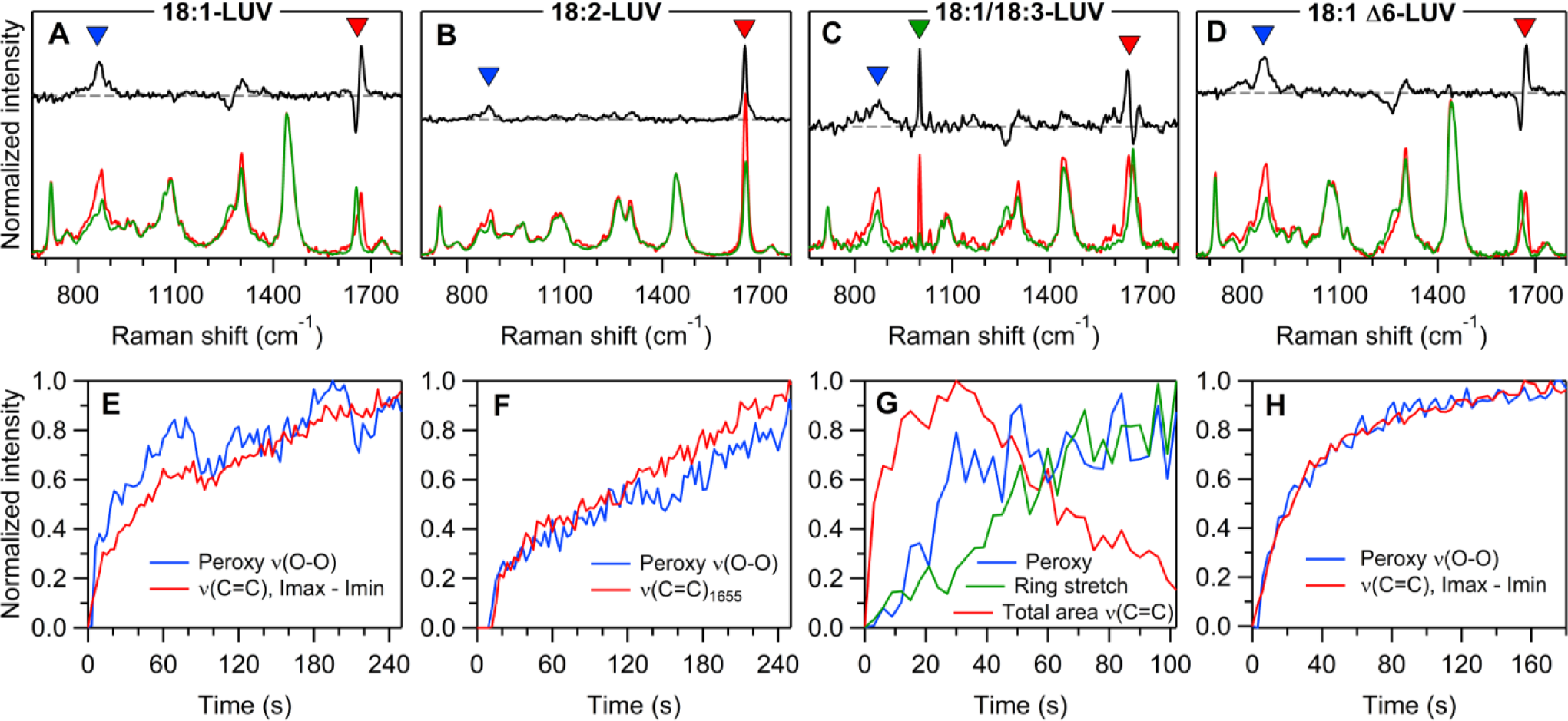
Chemical changes during vesicles oxidation. (A-D) Raman spectra corresponding to the beginning of the measurements (t=0 s, green curves), and to a specific characteristic time (red curves, A: 165 s, B: 216 s (B), C: 51s, D: 60 s). The 1:1 difference spectrum is in black. The characteristic band regions are tagged: peroxy (O-O) stretch (blue signs), carbon double-bond stretch (red signs) and 6-membered ring stretch (panel C, green signs). (E-H) Raman kinetics of characteristic band regions. The same colors than for tags in top panel are used: peroxy (O-O) stretch (blue curves), carbon double-bond stretch (red curves) and 6-membered ring stretch.

### Relation between chemical reactions and observed shape transitions

In presence of oxygen and under light activation, a photosensitizer can lead to two types of oxidative reactions induced by ROS. Type I oxidations are initiated by radicals and Type II oxidations by singlet oxygen. Because the Ce6 quantum yield is important, the latter process is expected to be dominant in our experiments (32). To determine the oxidative nature of the light-induced membrane responses and to discriminate which chemical mechanism was dominant, we used sodium azide as quencher of singlet oxygen. None of the GUVs formed in 50 mM sodium azide solution showed sign of oxidation upon illumination.

These observations are in agreement with our Raman *in situ* analysis, highlighting that the lipid tail C=C double bonds are the privileged site of the ROS reactions. The subsequent production of hydroperoxides is manifested by an increase of the Raman peak in 840-880 cm^-1^ region for all the types of vesicles except trans-DOPC and DMPC. The production of such a polar group within the hydrophobic area of the membrane strongly changes the structure of the lipid tail. As this group tends to migrate near the polar zone of the lipids head, this process tends to increase the membrane surface area, leading to the dynamics observed during Stage 1 (Fig. 3) (28, 29, 34). As expected, these two processes are very fast and temporally correlated for all lipids used in this study. Furthermore, we found that the variation of membrane surface area related to the hydroperoxidation depends on the position on the unsaturation, as seen from the large amplitude of the fluctuations during Stage 1 of 18:1∆6-GUVs photooxidation (Fig. 4). This can be explained by the fact that the 18:1∆6-GUVs double bound is positioned closer to the head group than for the other compositions, facilitating its relocation toward the lipid-water interface in the hydrophilic zone of the membrane (57).

During Stage 2 of the vesicles' photooxidation dynamics, various degradation products are produced (20, 21) and expelled from the hydrophobic core of the membrane because of their increased hydrophilicity. In consequence, such species are not present in the recorded Raman spectra of immobilized vesicles. However, the decrease in double bonds of the tails can be observed in some experiments. The cleaved lipids produced within the oxidized membrane are known to increase membrane permeability, as observed in our experiments during phase 2 (Fig. 4). They also weaken the membrane. Accordingly, at this stage, the optically trapped vesicles are destabilized and burst, leading to a loss of Raman signal. Moreover, cleaved lipids are known to increase membrane permeability (15, 23)

Throughout Stage 2 the oxidation of the membrane continues, as highlighted by the successive swell-burst cycles of the vesicles resulting from the accumulation of oxidation products. The membrane composition continually changes over the oxidation. So, it would be expected that the permeability of the membrane would increase with increasing oxidation degree. This is clearly not the case and, after an abrupt increase of *P_s_* corresponding to the initiation of Stage 2, *P_s_* remains constant (18:1- and 18:2-GUVs) or even slightly decreases (18:1/18:3-GUVs). This evidences that the membrane permeability is not directly and quantitatively related to the oxidation level and that more complex phenomenon have to be taken into consideration. For example, the asymmetric distribution of oxidative damages across the bilayer has been shown to affect membrane permeability (29, 36). Hence, structural or dynamics properties of the membrane can be modulated over oxidation (16, 53, 58), in addition to possible effects of cleaved tail fragments on the membrane properties (15).

## CONCLUSIONS

In this work, we studied the influence of lipid unsaturation on the dynamics evolution of the physical and chemical properties of GUVs undergoing photooxidation.

Tracking the morphology of the GUVs made of various unsaturated lipid compositions, we identified two consecutive stages upon illumination: a Stage 1 where the vesicles exhibit shape destabilizations followed by a Stage 2 associated with a rounding of the vesicles, decrease of their size and optical contrast and occurrence of swell-burst cycles. Combining quantifications of these dynamic morphological changes and theoretical analysis, we estimated the decay of membrane permeability to sucrose with time. We further showed that the GUVs lytic strain was decreased during the course of the illumination, pointing out to a weakening of photooxidized membranes. Interestingly, we evidenced a complete absence of oxidation of the trans fatty chains. We showed that the onset of photooxidation is independent of the number of lipid unsaturation (18:1- and 18:2-GUVs), but is faster if the position of the unsaturation is close to the head group (18:Δ6-GUVs). However, the rate of optical contrast decay – related to the permeabilization of the membrane – is increased by both the number of double bond and the proximity of the unsaturation to the head group. Interestingly, the permeabilization of the membrane is not proportional to its oxidation level and would be governed by a threshold effect.

In order to relate the dynamic morphological changes to the chemical processes occurring *in situ* during the photooxidation process, we conducted the RTM experiments on optically trapped LUV. These Raman measurements allowed us to observe directly the long-scale kinetics with time resolution of 3 seconds, of the following photoinduced processes: C=C bond isomerization, conjugation of C=C bonds, peroxide formation and a cyclization reaction involving the 6-membered ring structure.

Our findings shed light on the fundamental dynamic mechanisms underlying the oxidation of lipid membranes and highlight the role of unsaturations on their physical and chemical properties. This work has important implications in the understanding of organelles and cell membrane fate under oxidative stress and has the potential to help the development of preventive treatments against premature cell aging.

## ASSOCIATED CONTENT

### Supporting Information

Detailed RTS analysis

Theoretical analysis of the swell-burst cycles

## AUTHOR CONTRIBUTIONS

S.B. designed research; A.B., M.C. and S.G.K. performed research; M.C., P.R. and S.B. contributed analytic tools; A.B., S.G.K., N.P. and S.B. analyzed data; A.B., S.G.K., M.C., P.R., N.P. and S.B. wrote the manuscript.

### ACKNOWLEDGMENT

We thank C. Vever-Bizet and F. Sureau for helpful discussions. We are grateful for financial support by Region Ile-de-France in the framework DIM Nano-K (S.B. and A.B.). M.C. and P.R. acknowledge the support of ONR N00014-17-1- 2628 award.

