## Supporting Material for "Lipid unsaturation properties govern the sensitivity of membranes to photo-induced oxidative stress"

Aurélien Bour,[1] Sergei G. Kruglik,[1] Morgan Chabanon,[2] Padmini Rangamani,[2] Nicolas Puff,[3,4] and Stéphanie Bonneau[1,*]

[1] Sorbonne Université, Faculté des Sciences et Ingénierie, CNRS, Laboratoire Jean Perrin, UMR 8237, 4 place Jussieu, Paris, France.

[2] Department of Mechanical and Aerospace Engineering, University of California San Diego, La Jolla, CA 92093, United States

[3] Sorbonne Université, Faculté des Sciences et Ingénierie, UFR 925, 4 place Jussieu, Paris, France.

[4] Univ Paris Diderot, Sorbonne Paris Cité, CNRS, Laboratoire Matière et Systèmes Complexes (MSC), UMR 7057, 75013, Paris, France.

### S1. THEORETICAL ANALYSIS OF THE SWELL-BURST CYCLES

In this section we aim to derive mathematical tools to gain information about the membrane properties based on the measurement of the GUVs contrast decay and radius dynamics. The theoretical framework of this model is based on Chabanon et al. (1).

#### S1.1. Evaluation of the membrane permeability to sucrose

We consider a spherical vesicle of volume $V=4/3\pi R^3$ with an inner sucrose concentration $c_s$. The total amount of sucrose in the vesicle is therefore $Vc_s$, and varies through (i) permeation through the membrane with a permeability to sucrose $P_s$, and (ii) by diffusion/advection through the pore of area $\pi r^2$ when it is open. Therefore the evolution of sucrose concentration in the vesicle can be expressed as

$$\frac{d}{dt}(V c_s) = -\pi r^2 \left( D_s \frac{\Delta c_s}{R_0} + v_L c_s \right) - A P_s \Delta c_s \quad , \qquad \text{(Eq. S1)}$$

where $D_s$ is the sucrose diffusion coefficient in water, $\Delta c_s$ is the sucrose concentration differential, $v_L$ is the leak-out velocity through the pore, and $A= 4\pi R^2$ is the membrane surface area. Let us consider one particular swell-burst cycle, say cycle number $n$. Since the pore lifetimes (~0.05 s) is much smaller than the cycles periods (~10 s), we assume that the cycle period $\Delta t_n$ is equal to the time required for the vesicle to swell from the resting radius $R_{0n}$ to $R_{fn}$, the radius at which the membrane ruptures. Here, subscripts $n$ denotes the cycle number. During that time, the pore is closed so the first term proportional to the pore radius $r$ in Equation S1 is null. Furthermore, the surrounding bath volume being much larger that the vesicle volume, we assume the external sucrose concentration to remain zero at all time. Developing the left hand side of Eq. S1 such as

$$\frac{d}{dt}(V c_s)=\frac{4}{3}\pi R^3 \frac{d c_s}{dt}+c_s 4\pi R^2 \frac{dR}{dt}\simeq\frac{4}{3}\pi R^3 \frac{d c_s}{dt}+c_s 4\pi R^2 \frac{\Delta R_n}{\Delta t_n} \quad , \qquad \text{(Eq. S2)}$$

where $\Delta R_n = R_{fn}$- $R_{0n}$, and $\Delta t_n$ is the period of the cycle n, we can rewrite Eq. S1 for the n^th cycle such as

$$\frac{d c_s}{dt}\simeq\frac{-3}{R_{0n}}\left(P_s+\frac{\Delta R_n}{\Delta t_n}\right)c_s \quad , \qquad \text{(Eq. S3)}$$

where $R_{0n}$ is the initial vesicle radius of the n$^{th}$ cycle. Assuming that the membrane permeability to sucrose is constant within one cycle, we can define the (constant) characteristic time

$$\tau=1/\left[3\left(\frac{P_s}{R_0}+\frac{\Delta R_n}{R_0 \Delta t_n}\right)\right] \quad , \qquad \text{(Eq. S4)}$$

so that the sucrose concentration satisfying Eq. S3 follows an exponential decay such as

$$c_s(t)=c_s(t_{0n})e^{-(t-t_{0n})/\tau} \quad , \qquad \text{(Eq. S5)}$$

where $t_{0n}$ is the time at which the cycle starts. Given that $\Delta R_n$, $\Delta t_n$, $R_{0n}$, and $t_{0n}$ are constant quantities that can be measured for each cycle, Eq. S4 can be fitted to each cycle to determine τ, and therefore the membrane permeability to sucrose $P_s$.

**S1.2 Relation to osmotic differential in oxidative products**

Given that sucrose and glucose are initially at equimolar concentration and are permeating through the membrane at a similar rate, they do not contribute to the osmolarity imbalance. Therefore the only solutes responsible for the osmotic swelling of the GUVs are the products released during the oxidation process. Let us denote $\Delta c_p$ the concentration difference of oxidative products across the membrane responsible for the osmotic imbalance. The volume conservation of the vesicle can be written as

$$\frac{dV}{dt}=A\frac{P_w \nu_w}{k_B T N_A}\left(k_B T N_A \Delta c_p-\Delta p_{Lap}\right)-\pi r^2 v_L \quad , \qquad \text{(Eq. S6)}$$

where $P_w$ the membrane permeability to water, $\Delta p_{Lap} = 2\sigma/R$ is the Laplace pressure with $\sigma$ being the membrane tension, $\nu_w$ is the molar volume of water, and $k_B$, $T$, and $N_A$ are the Boltzmann constant, the bath temperature, and the Avogadro's number respectively.

Once again, we consider the evolution of the vesicle volume during the swelling stage of a cycle number $n$. Furthermore, we assume that the osmotic pressure dominates the swelling phase ($k_B T N_A \Delta c_p >> \Delta p_{Lap}$) so that Laplace pressure is neglected. This assumption is supported by the fact that the radii increase are quasi-linear (see Figure 3). We get

$$\frac{dR}{dt}=P_w \nu_w \Delta c_p \quad . \qquad \text{(Eq. S7)}$$

Assuming the oxidative product concentration increases linearly during the time of a cycle, we have

$$\int_0^{\Delta t} \Delta c_p \, dt=\int_0^{\Delta t} \left(\alpha_n t+\Delta c_{pn0}\right) dt=\frac{\alpha_n}{2}\Delta t^2+\Delta c_{pn0}\Delta t \quad , \qquad \text{(Eq. S8)}$$

where is the production rate of oxidative products (mol/m3/s), and is the initial oxidative product concentration of cycle number n. We therefore deduce from Eqs. S7 and S8

$$\frac{\Delta R_n}{\Delta t_n} = P_w \nu_w \left( \frac{\alpha_n}{2} \Delta t + \Delta c_{pn0} \right) \quad . \qquad \text{(Eq. S9)}$$

Our measures show that $\Delta R_n/\Delta t_n$ decreases with increasing cycle numbers. Two scenarios can be considered: (i) the production of oxidative products happens mainly at early time and is then null for most of the process ($\alpha_n$=0). In that case $\Delta c_{pn0}$ decreases at each circle by leak-out through the transient pore, in a process similar to the osmotic swell-burst cycle described in Chabanon et al (1). Assuming $P_w\nu_w$ constant, the drop in oxidative product at each cycle can therefore be estimated from the measure of $\Delta R_{n+1}/\Delta t_{n+1}$ - $\Delta R_n/\Delta t_n$. In scenario (ii), oxidative products are released during the course of the swell-burst cycles ($\alpha_n$>0). Given the Raman kinetics, it is likely that the release rate $\alpha_n$ decreases at each cycle. Note that because $\alpha_n \Delta t_n$ is positive, this second scenario still implies that $\Delta c_{pn0}$ decreases at each cycle by leak-out through the pore.

### S2. DETAILED RTM ANALYSIS

| Frequency range | Assignment |
|---|---|
| 1720 – 1750 | ν(C=O) in ester COOR |
| 1665 – 1675 | $\nu(C=C)_{trans}$ in acyl chain |
| 1650 – 1660 | $\nu(C=C)_{cis}$ in acyl chain |
| 1430 – 1470 | Scissoring $\delta(CH_2, CH_3)$ in acyl chain |
| 1295 – 1305 | twisting $\delta(CH_2)$ in acyl chain |
| 1260 – 1270 | rocking $\delta(=CH_2)_{cis}$ in acyl chain |
| 1050 – 1180 | ν(C-C) in acyl chain |
| 950 – 1000 | $\nu_{as}(O\text{-}C\text{-}C\text{-}N^+)$, wagging $\delta(=CH_2)$ |
| 820 – 900 | $\nu_s(O\text{-}C\text{-}C\text{-}N^+)$, $\nu_{as}(C_4\text{-}N^+)$, rocking $\delta(CH_2)$ |
| 840 – 880 | peroxide ν(O-O)[a] |
| 717 | $\nu_s(C\text{-}N^+)$ of polar head |

**Table S1.** Frequencies ($cm^{-1}$) and assignment (2, 3) of the major bands in Raman spectra of lipid vesicles studied. ν = stretching mode (s, symmetric; as, asymmetric). δ = deformation bending mode (in-plane: scissoring or rocking; out-of-plane: twisting or wagging). [a]Assignment of transient peroxide band region according to Vacque & al. (4).

#### S2.1. DOPC vesicles

In the case of DOPC, the most pronounced spectral changes occur in three spectral regions associated with: (i) carbon double-bond stretching ν(C=C), in 1650-1675 $cm^{-1}$; (ii) rocking deformation $\delta(=CH_2)$ at 1265 $cm^{-1}$, also involving the carbon double bond; and (iii) peroxide ν(O-O) stretch (4), which appears in difference Raman spectra within 840-880 $cm^{-1}$. The observed spectral changes reveal two major photo-induced processes that occur in lipid membrane after Ce6 photo-activation: *cis-trans* isomerization and peroxide formation.

*Cis-trans* isomerization is characterized by the decrease of $\nu(C=C)_{cis}$ band intensity at 1655 $cm^{-1}$ with simultaneous increase of the $\nu(C=C)_{trans}$ band intensity at 1670 $cm^{-1}$ which is a characteristic frequency of the *trans*-species (5). The corresponding difference Raman spectra contain characteristic, rather symmetric dispersive contour with the minimum at 1655 $cm^{-1}$ and the maximum at 1670 $cm^{-1}$; this contour becomes more and more pronounced with the lapse of time. The appearance of *trans*-DOPC species occurs immediately (within 3 s time resolution of our experiment) after photo-excitation. It takes about 54 s for $\nu(C{=}C)_{trans}$ band to become equal in intensity with the $\nu(C{=}C)_{cis}$ band. It is important to note that the total Raman intensity within the range 1600-1700 $cm^{-1}$, containing ν(C=C) stretch of both *cis*- and *trans*-DOPC, remains roughly the same with a weak tendency to the increase with time, revealing that no other, besides these two, major transient structural species is involved. The Raman intensity of rocking deformation $\delta(=CH_2)_{cis}$ in the 1260-1270 $cm^{-1}$ range experiences kinetic changes very similar to those of $\nu(C{=}C)_{cis}$, but with a decreased magnitude.

Peroxide formation in DOPC vesicles has been revealed by substantial increase of Raman intensity in the characteristic ν(O-O) frequency range 840-880 $cm^{-1}$ (4). Since this spectral region is congested by numerous Raman bands (Table S2), the kinetic changes have been analyzed from difference Raman spectra obtained by subtraction of normalized Raman spectra of vesicles with and without addition of Ce6. Peroxide species appear immediately (within our 3 s time resolution) after photo-excitation with very high amplitude (~35%) suggesting that the process of peroxide formation is very efficient from the very beginning.

The influence of the carbon double-bond position on lipids oxidation has been studied for the vesicles consisting of DOPCΔ6. The general picture of Raman intensity changes was found to be quite similar to the previous case of DOPC; however some notable differences were observed as well. In particular, the process of *cis-trans* isomerization proceeds much faster in DOPCΔ6: it takes ~20 seconds for $\nu(C{=}C)_{trans}$ band to become equal in intensity with the $\nu(C{=}C)_{cis}$ band, versus ~54 seconds for DOPC. At the same time, it is interesting to note that the fast components of peroxide kinetics are rather similar for DOPC and DOPCΔ6: in both cases it takes about 20 seconds to reach the 50% level of peroxide band intensity.

Another very interesting observation consists in the following: no photo-induced changes in Raman spectra of vesicles consisting of DOPC*trans* were observed at any time delay after irradiation beginning. This finding suggests that the peroxide formation requires *cis*-configuration of the carbon double bond in monoene lipids in the presence of activated Ce6.

#### S2.2. DLPC vesicles

For DLPC vesicles, in addition to peroxide formation, we observe increase of C=C stretching band instead of *cis-trans* isomerization. Indeed, the $\nu(C{=}C)_{cis}$ band at 1657 $cm^{-1}$ experiences strong intensity enhancement with the lapse of time, along with the small frequency downshift to 1654 $cm^{-1}$. It is known that, in normal conditions, there exists a linear relationship between the number of C=C bonds and the intensity ratio $R=I(\nu_{1667})/I(\delta_{1444})$ (2, 6). After normalization of this ratio R to 2 double bonds in the case of DLPC without Ce6, from normalized Raman spectra at different time delays, we obtained R=3 at $\Delta t=80$ s and even R~3.7 at $\Delta t=280$ s, with the kinetics being stabilized around this value for longer time delays. Thus, our Raman data suggest that Ce6 photosensitizer activation causes the increase of the number of C=C double bonds in linoleic chain, or, more probably their delocalization and conjugation, in DLPC-vesicles. At this early stage, the intensity of the band at 1264 $cm^{-1}$ remains unchanged with respect to that of non-conjugated DLPC (5, 7).

It should be noted that in some experiments (data, not shown), when the experiment was extended to longer time delays, we eventually observed the reverse process of moderate intensity decrease of the $\nu(C{=}C)_{cis}$ stretch at 1654 $cm^{-1}$ with concomitant appearance of weak *trans*-species manifested by a shoulder at 1670 $cm^{-1}$ and the intensity decrease of the $\delta(=CH_2)_{cis}$ deformation band at 1264 $cm^{-1}$.

The initial peroxide formation for DLPC vesicles (Fig. 5F) is again very rapid, within 1-2 time points (3-6 seconds), although being less efficient (<20%) than in the case of monoenes (30-35%, Fig. 5B,D). In the case of

DLPC vesicles, it takes more than 150 seconds to reach the 50% level of peroxide band maximal intensity (Fig. 5F) in contrast to ~20 seconds in the case of DOPC vesicles (Fig. 5B).

#### S2.3. DOPC/DLPC vesicles

Finally, in the case of the vesicles containing lipids mixture 18:3/18:1, the observed spectral changes and kinetics are rather complicated. In addition to a superposition of photoinduced processes observed for DOPC and DLPC, namely *cis-trans* isomerization, C=C double bond delocalization and conjugation, and peroxide formation, we observe also a cyclization reaction. Indeed, the creation of a six-membered (toluene-like) ring is manifested by the appearance of strong and narrow characteristic Raman bands at 1000, 1031, and 1596 $cm^{-1}$. Moreover, the kinetics of photoinduced changes vary from one measurement to the other reflecting the competition of simultaneously occurring processes; one set of the observed kinetics is shown in Fig. 6.
